# High Throughput Detection of Capillary Stalling Events with Bessel Beam Two-Photon Microscopy

**DOI:** 10.1101/2022.12.16.520779

**Authors:** John Giblin, Sreekanth Kura, Juan Luis Ugarte Nunuez, Juncheng Zhang, Gulce Kureli, John Jiang, David A. Boas, Ichun A. Chen

**Affiliations:** Boston University Department of Biomedical Engineering Boston, MA; Boston University Neuorophotonics Center Boston, MA

## Abstract

Disruptions in capillary flow have the potential to drive pathology across numerous diseases. But our understanding of the temporal and spatial dynamics of these events are hindered by slow volumetric imaging rates and the reliance on laborious manual analysis to process data. To address the challenges of increasing volumetric imaging speed, we use a custom-built Bessel beam two-photon microscope for efficient volumetric imaging of the capillary network. We demonstrate its ability to continuously monitor roughly 200 capillaries for capillary flow stoppages (i.e. stalling events) at a frame rate of approximately 0.5 Hz and develop a semi-automated correlation-based approach for identifying these stalling events. We applied our system and algorithm in a photothrombotic model of stroke and show elevated levels of stalling 1-week post-stroke in regions both within and outside of the stroke region, demonstrating that stalling may have impacts on stroke recovery that extend past the acute stage.

## Introduction

Adequate blood flow is essential to supporting healthy brain function^1^. Global and regional reductions^2–6^ in cerebral blood flow as well as alterations in hemodynamic^3,6^ responses are associated with or precede cognitive decline across multiple neurodegenerative diseases including Alzheimer and Parkinson’s. At the smallest scale, alterations in microvascular structure^7^ and flow can also drive pathology^8–10^, including regions affected by stroke induced hypoxia. This includes transient disruptions in flow, or “stalls”, in normally continuous cerebral capillary flow^11–13^. Recent work has demonstrated the potential role of stalls in pathology including Alzheimer’s disease^14^ and stroke^15–18^. In addition to potentially inducing local hypoxia, these events can contribute to total vascular resistance^14^, as well as flow heterogeneity^19^, further inducing hypoxia^20^. Treatments to reduce this stalling have been associated with favorable cognitive outcomes in a model of Alzheimers^14^ and reduced infarct size in stroke^15,18^.

Investigations into the effects of stalling and its potential as a therapeutic target have been hindered due to technological limitations. Traditional two photon imaging z-stacks are frequently used to measure stalling rates^12,14–17^, but only observe individual capillaries for a small fraction of the acquisition time. Further, even when a stall is observed, the onset and cessation of events is typically not seen. The alternative is to monitor a single XY plane over a long duration, but due to two photon confinement, only a handful of capillaries are typically visible in a single plane. Optical Coherence Tomography (OCT) angiography provides continuous volumetric monitoring of capillaries^13,18^ but lacks the temporal resolution to catch shorter stalling events. In addition, OCT also lacks the ability to perform multicolor fluorescence imaging like two-photon microscopy, which has aided in investigating the source of the stalling events^14,15,17,18^.

In addition to these limitations, analyzing and extracting stalling information for these enormous datasets is highly labor intensive. For both two-photon microscopy and OCT, images must be manually analyzed to identify the stalling events. In the case of two-photon microscopy, analysis must be crowd sourced to citizen scientists who volunteer to sift through hundreds of thousands of capillary images to identify stalled capillaries so that the resulting statistics can be reported in a reasonable amount of time^14,21^.

Further, the sparse nature of stalling in these data makes the construction of a large training dataset for automation difficult due to class imbalance. As more complex investigations are performed, and therapeutic modulation of stalling continues to be explored, the amount of data that needs analysis will grow significantly. Therefore, there is a need for high throughput imaging and analysis of stalling parameters to avoid a data processing bottleneck.

To address these challenges, we use a custom Bessel beam two photon microscope^22^ to identify stalling events by continuously monitoring a 713×713×120 μm volume acquired at 0.57 Hz, roughly four times the volumetric imaging rate of OCT angiograms. We then developed a correlation-based detection approach to semi-automate the analysis for stall identification, we show a reduction of 50% in the time required to analyze the data and a 2 to 3 times improvement in the number of stalls identified when compared with unassisted analysis. With the combined improvements in temporal resolution and improved detection rate, we found that a majority of stalls are very short in duration, making it likely they were missed in other studies. This enabled us to observe differences in stall dynamics between young and aged mice, as well as track changes in stall dynamics before and after stroke in the same capillary network.

## Results

Fluorescence images acquired using a two photon microscope with an axicon based Bessel module having an extend foci full width half max of 100 μm is shown in (Figure 1a). Within this volume, roughly 200 capillaries are monitored for both short and long duration stalling events (Figure 1b; Supplemental Movie 1), where red blood cell shadows (RBCs) are stationary for at least two or more consecutive frames. Identification of stalled capillaries, and the times they were stalled are used to calculate key stall parameters^13^ (Figure 1c). This includes the percent of all capillaries that undergo stalling at any time (incidence), the average percent of capillaries actively stalling at a given time (point prevalence), and the average total time a given capillary spends stalled during the measurement duration (cumulative stall duration). Identification of stalling capillaries and annotating the time and duration of each stall event for the acquired time series data is a particularly laborious task since there are hundreds of capillaries to observe at higher volume rates than previous studies. To make stalls more apparent, the data is broken down into sub-ROIs containing approximately 20-30 capillaries each, which is then scrolled through frame by frame to identify instances where are all RBC shadows within a capillary segment are stationary between two or more frames. This process is repeated for all sub-ROIs until the whole field of view has been processed. A full 713×713×120 μm dataset of 350 frames take close to 12 hours per dataset and the results can be variable depending on the focus and fatigue level of the observer. To improve both throughput and detection accuracy, we introduced a metric that can predict the occurrence of a stall in each capillary.

**Figure 1.**
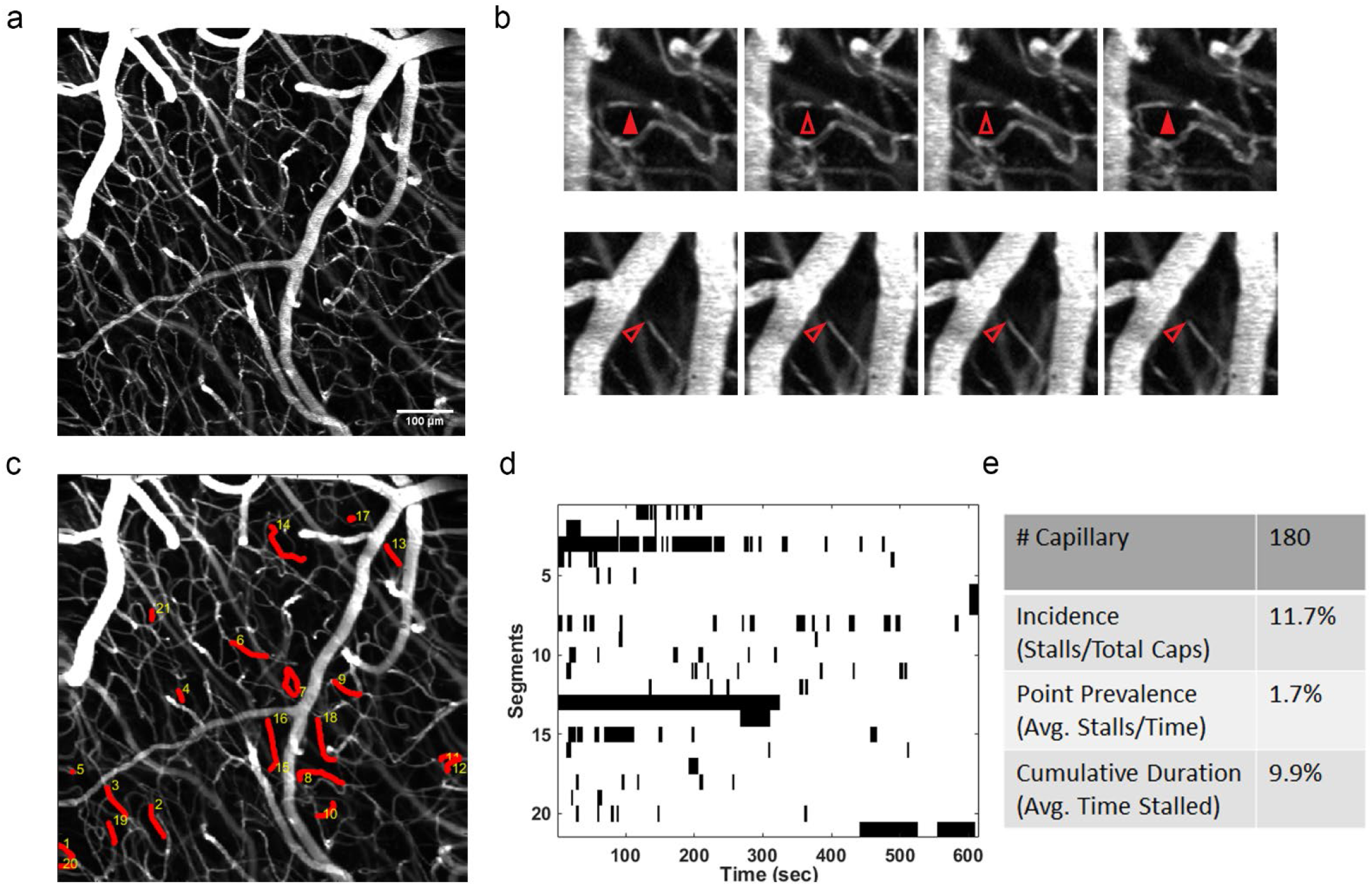
(a) MIP of a 713×713×120 μm volume scanned at 0.57 Hz for 10 minutes to observe instances of capillary stalling (b) Representative examples of types at stalling events captured during imaging. (Top) Red blood cell (RBC) shadows are stationary for 2 sequential frames during a short stalling event before normal flow resumes. (Bottom) A shadow is continuously present in the same position along the length of the capillary during a long stalling event. (c) MIP with stalled capillaries marked in red (d) Stallogram show the times each capillary was stalled for (marked in black) (e) Stall statistics calculated based on the stallogram and total number of capillaries.

During normal flow, at the scanning rate used, new shadows (mostly RBCs) will be present in every frame, varying in number and position. During a stall, the same shadows will remain in the same position along the length of the capillary until flow resumes in that capillary. Therefore, the intensity along the length of the vessel is expected to be well correlated during stalling events. To facilitate extraction of the intensity along the length of the capillary, we semi-automatically extract the centerline trace of capillaries of interest ^23^(Figure 2a). This generates images of intensity along the capillary length versus time (LT images). Plotting the frame-to-frame intensity correlation versus time shows a clear increase during stalling events, before returning to lower levels (Figure 2b). To validate this metric, we threshold the correlation and use it as a predictor of stalling, which was then confirmed by a human observer.

**Figure 2.**
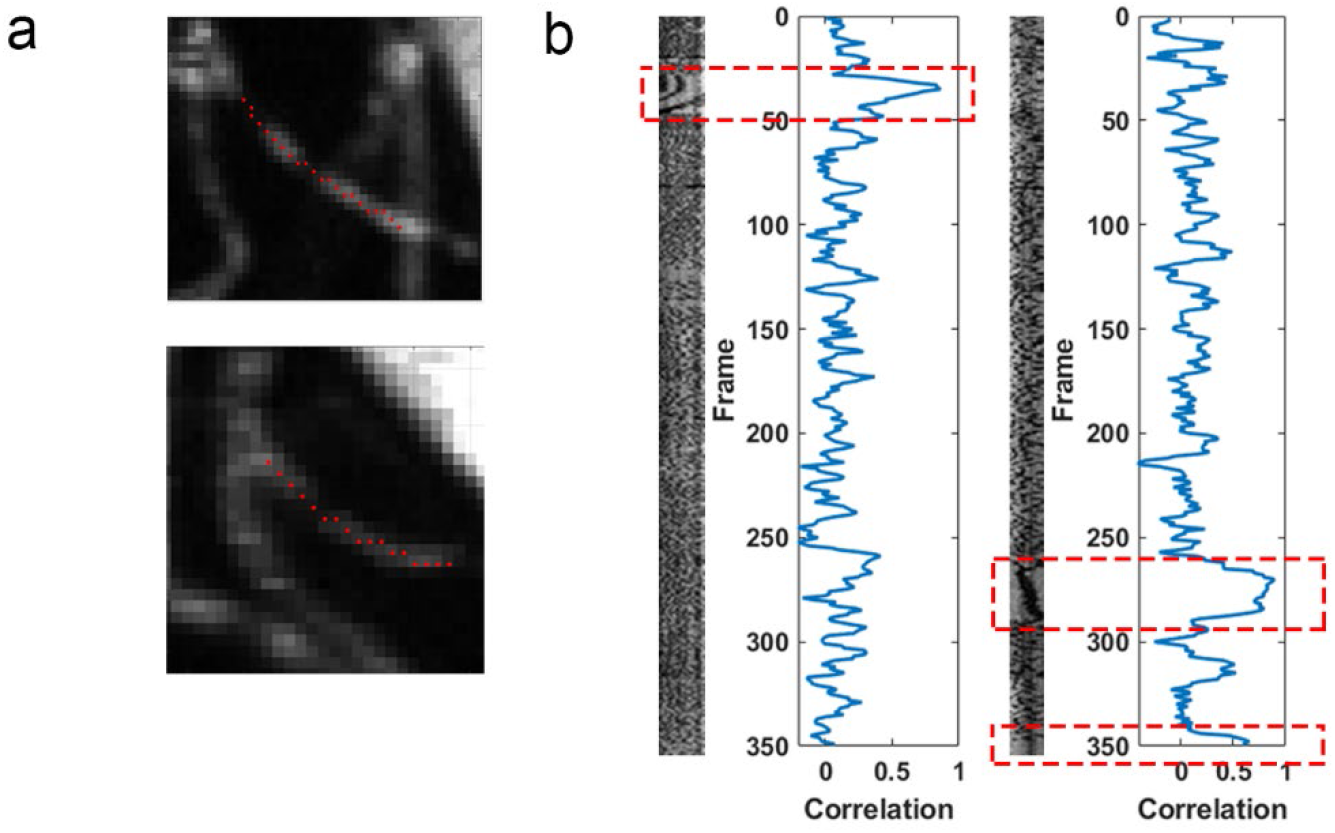
(a) Capillaries that undergo stalling with the centerline drawn automatically after single click selection (b) Length versus time plots of the capillaries from (a) with the correlation versus time plotted alongside. Red boxes indicate times where the capillary was stalled and the same RBCs are present for multiple frames, which corresponds to a rise in frame-to-frame correlation.

### Correlation Validation

To validate our approach, we first manually analyzed 10 datasets from 4 mice (4-8 Months old). For each dataset we then used our correlation-based approach to predict stalls from length versus time (LT) images generated for each capillary. The threshold was set intentionally low, trading off specificity for increased sensitivity. The result was then analyzed by another observer to determine if flagged events were a stall (true positive) or not (false positive) (Table 1). This thresholded correlation generated significantly more false positives than true positives, meaning that it could not be directly used for calculation of stall statistics. However, while this correlation-based identification approach was unable to reliably predict stall parameters accurately, we found that the validation process was roughly twice as fast as the original blind manual analysis (going from 10-12 hours to about 6 hours per ROI). It could therefore be used to improve analysis throughput. In addition, we found 2 to 3 times as many stalling events with this new approach indicating that the manual process alone had a large number of false negatives. Further, the few false negatives for the correlation-based approach (stalls found manually by the observer but not flagged by correlation) gives confidence that we are capturing as many stalling events as possible and that we don’t need to confirm capillaries marked as negative by the correlation-based approach, further speeding the manual validation process. This correlation-based identification approach can therefore be used as an analysis aid that can speed up the analysis process while providing more precise estimation of the stalling statistics.

**Table 1.** Truth table of capillaries that stall found via manual inspection of the data (left) and correlation thresholding of the LT plot (right). A separate observer made ground truth determination in all cases.

|  | Manual<br>Negative | Manual<br>Positive | Correlation<br>Negative | Correlation<br>Positive |
| --- | --- | --- | --- | --- |
| Ground Truth<br>Negative | 2074 | 37 | 961 | 1150 |
| Ground Truth<br>Positive | 245 | 102 | 30 | 217 |

### Comparison to OCT Stalling

We then compared our stall parameters calculated from the results of the correlation aided detection to stalling statistics previously reported with OCT (Table 2)^13^. We found stalling incidence was twice that reported by OCT, likely due to the increased temporal resolution of the Bessel beam microscope. When looking at the distribution of the duration of the stalling events (Figure 3d), most of the events were less than 8 seconds in duration and therefore could easily be missed by the prior OCT angiography results which took 9 seconds to repeat frame imaging. This also contributes to the lower cumulative stall duration we measured with the Bessel beam microscope, as OCT was only sensitive to longer stalls and thus does not identify many capillaries that have only short duration stalling events (as suggested by Figure 3a). To confirm this, we removed shorter stalls in our Bessel beam results and then recalculated the statistics. When we only analyze stalls occurring for ~8 sec or longer, our measured incidence is very similar to what was found previously with OCT (Figure 3a). The cumulative stall duration (Figure 3c) did remain lower, still likely due to the increased temporal resolution, and improved ability to identify the end of longer stalling events with the Bessel beam approach. The variability in stall point prevalence found in OCT (Figure 3b) was too high to make a precise comparison to that found with the Bessel beam results, but the results are none-the-less consistent.

**Figure 3.**
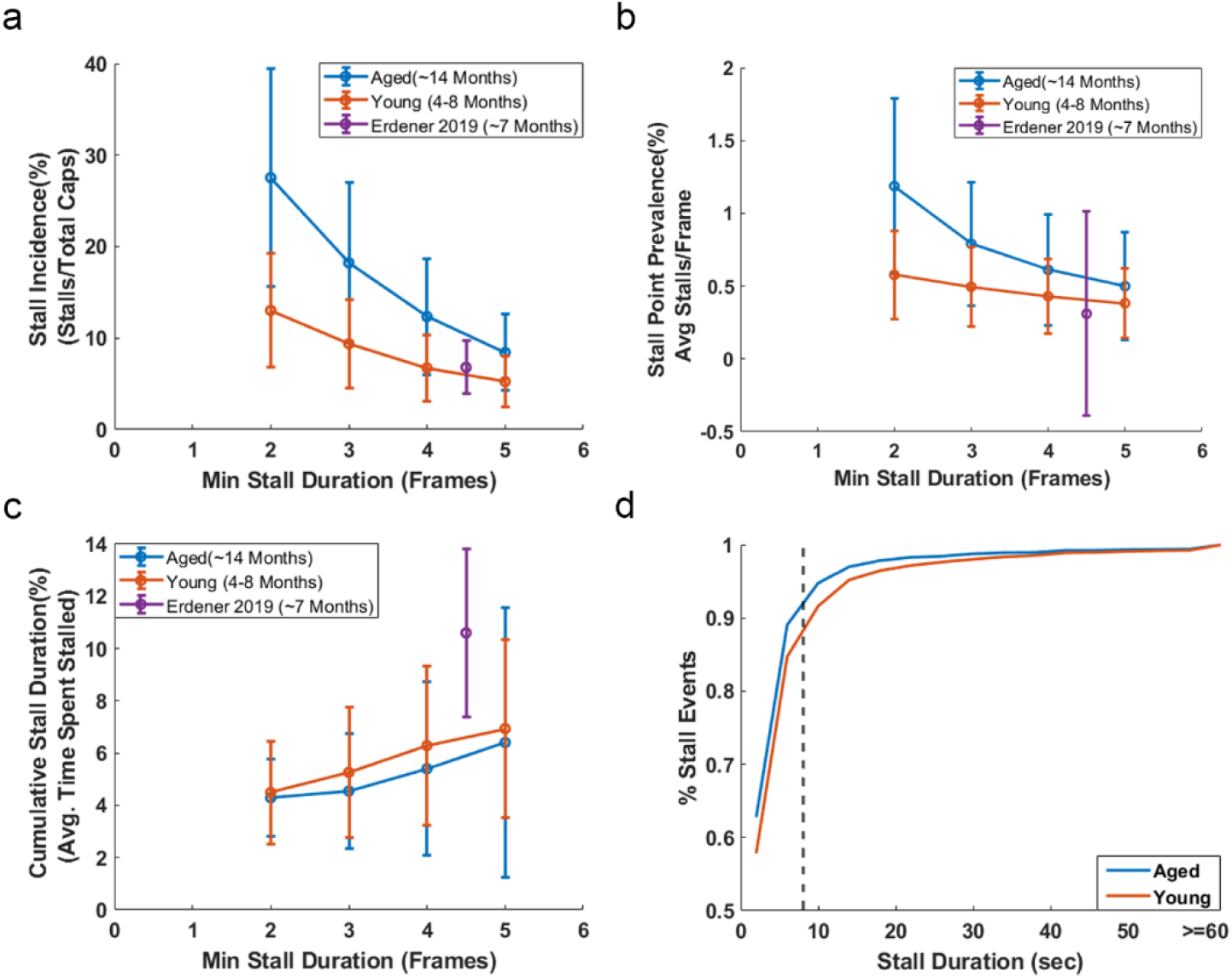
(a-c) Stalling statistics calculated after removing stalls shorter than a given length. Stall length of two includes all stalls detected. Purple point indicates the same statistics reported previously by OCT for a chronic window^13^ used to compare (a) stall incidence, (b) point prevalence and (c) cumulative stall duration. (d) Cumulative distribution function of all individual stalling events of young and aged groups. Dashed line represents the time to generate a single OCT angiogram.

**Table 2.** Comparison of stalling statistics measured by OCT (reported in Erdener et al 2019) and by Bessel beam two photon microscopy

|  | **OCT<sup>13</sup>**<br>**(~7 Months)** | **Bessel Beam TPM**<br>**(4-8 Months)** | **Bessel Beam TPM**<br>**(14 Months)** |
| --- | --- | --- | --- |
| ROIs | 12 | 10 | 11 |
| Frame Period | ~8 s | 1.75 s | 1.75s |
| Incidence | 6.8% +/- 2.9 | 13.02%+/- 6.2 | 27.5%+/-12 |
| Point Prevalence | 0.31% +/- 0.7 | 0.58%+/-0.3 | 1.18%+/-0.6 |
| Cumulative Stall Duration | 10.6% +/- 3.2 | 4.49%+/-1.97 | 4.29%+/-1.49 |

We applied this correlation aided approach to analyze data taken from older (n=4 mice,14-Month-old) mice and compared it to the results from the younger group analyzed in the validation study. The older cohort had increased rates of stalling compared to the younger group, predominantly of shorter duration. When only looking at longer duration stalls (4 frames or more, i.e., more than ~8 sec), stall incidence was similar to the young cohort and previous OCT results.

### Photothrombotic Stroke

To demonstrate the ability of our system to track changes in stalling statistics. We used a targeted photothrombotic model of stroke^24^ and measured the differences in capillary stalling at baseline compared with 1 week post stroke in 14 month old C57BL/6J mice. Regions-of-interest (ROIs) were determined to be in the stroke core, peri-infarct, and contra-lesional hemisphere as identified using SFDI to track changes in scattering (Figure 4b)^25^. In total there were 2 ROIs in the stroke core, 3 in the peri-infarct region, and 3 in the contra-lesional hemisphere. Both ROIs in the stroke core and all but one ROI in the peri-infarct region saw increased stalling incidence. The one ROI that did not show an increased incidence, had the highest incidence at baseline. We also found longer stall durations in all ROIs, including the ROIs in the contra-lesional hemisphere. The stalling point prevalence was increased, primarily driven by stalling duration, as it increased even in regions where overall incidence was unchanged. This shift is also apparent when looking at the distribution of the durations of the stalling events across all capillaries (Figure 4 f). At baseline, the older mice had higher incidence which was primarily driven by shorter stalling events. One week after stroke, longer duration events account for a larger percent of all stalls, even in the contra-lesional control ROI (Figure 4g).

**Figure 4.**
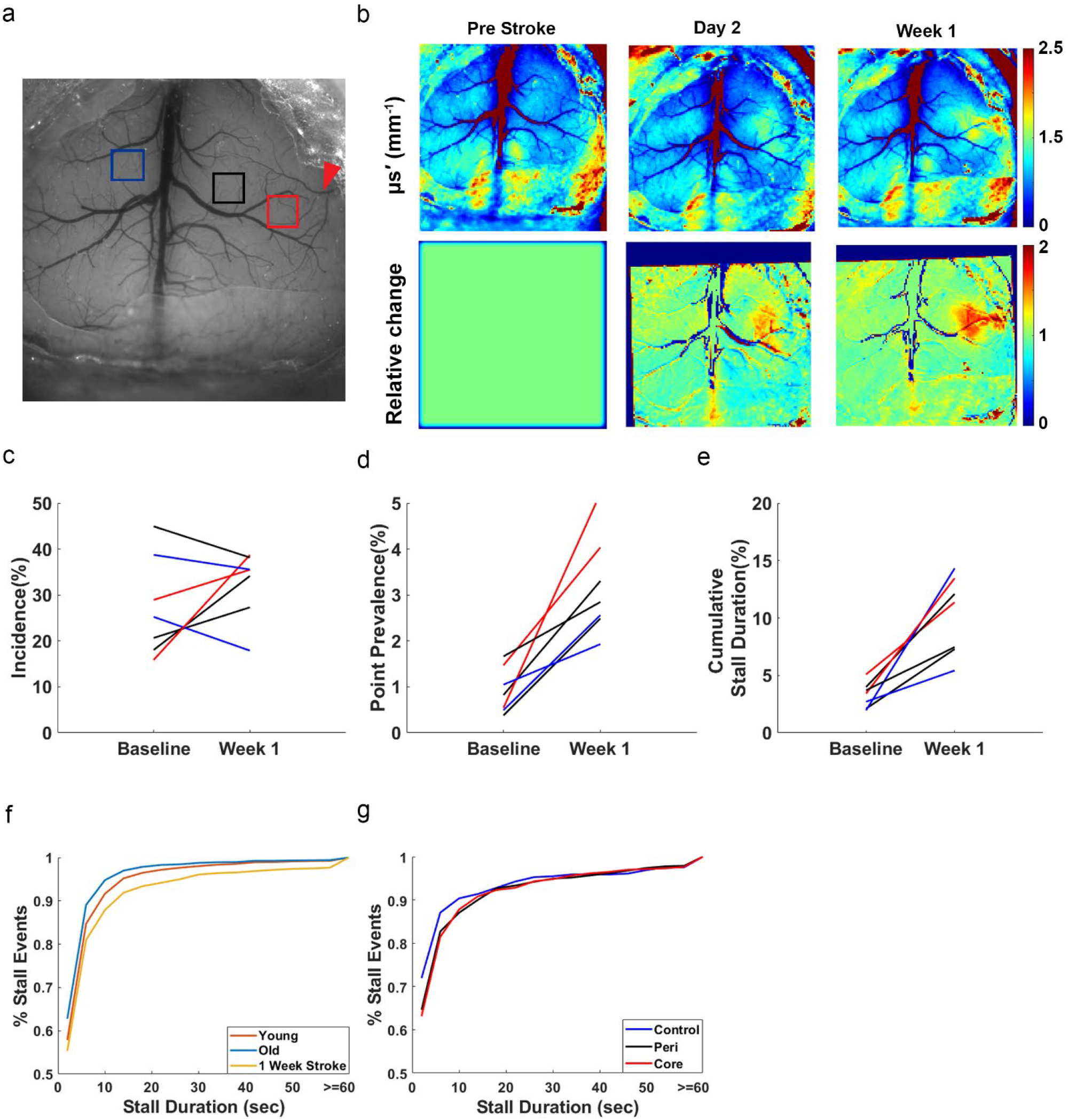
(a) Widefield reflectance image showing cortical vasculature at baseline. Red arrow indicates the artery that was targeted for photothrombotic stroke. Boxes show the ROIs that were imaged at baseline and 1 week post stroke. Colors indicate the relation of the ROI to the stroke. Red indicates stroke core, black peri-infarct and blue contra-lesional hemisphere. (b) SFDI scattering and relative images show the increase in scattering as a result of the stroke, used to determine the stroke core. (c-e) Changes in stall statistics in 7 ROIs across 2 mice from baseline to 1 week after stroke. Color indicates the relation to the stroke core. (f) Cumulative percentage of stalling events across all stalling capillaries as a function of duration across different groups. Line indicates the percentage of stalls with a duration less than or equal to the duration. (g) Cumulative percentage of stalling events across stalling capillaries based on region at 1-week post stroke.

### Arterial Dilations Transiently Reduce Stalls

In addition to volumetric imaging offered by the extended depth of field, Bessel beam allows for simple estimation of vessel diameter by using the vessel fluorescent intensity^22^ (Figure 5b). We hypothesized that periods of increased flow may reduce stalls by providing sufficient pressure across the capillary to clear the stall. We therefore analyzed the average stalling point prevalence around large, isolated dilation events (see Methods). We found that the average point prevalence was significantly reduced by the time of peak dilation (Figure 5c) and shortly after. This decline started prior to the peak diameter because flow starts to increase as soon as vessel diameter starts to increase.

**Figure 5.**
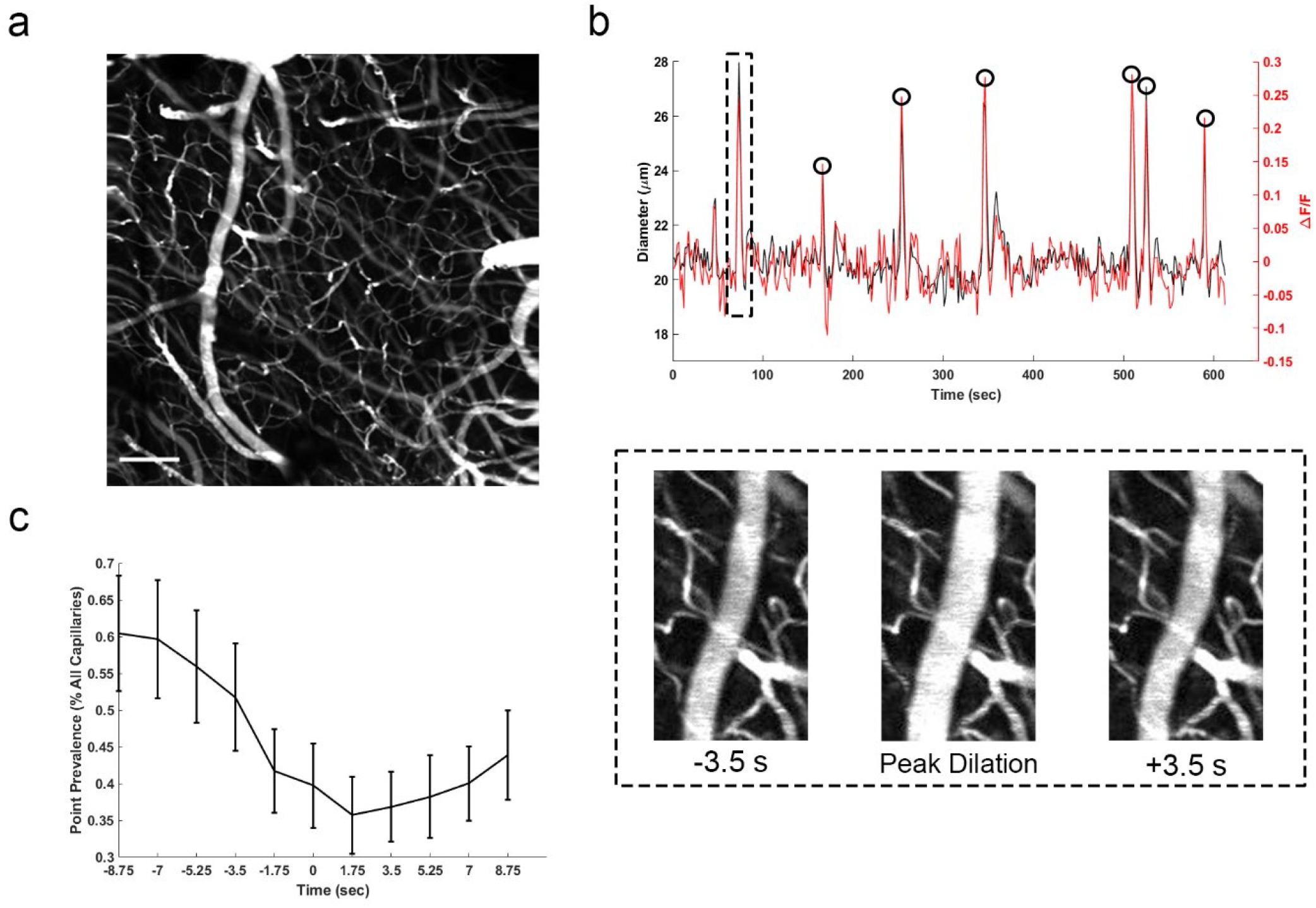
(a) Full stalling field of view with an artery on the left-hand side. Scale bar 100 μm (b) Diameter and fluorescence intensity trace of a section of the artery shows close correlation between fluorescence and diameter with the Bessel beam (Correlation coefficient: 0.88). Black circles indicate time points used for analysis in (c). (c) Changes in stall point prevalence relative to large dilation events across 9 ROIs. Time 0 corresponds to the time of peak diameter during dilation.

## Discussion

To overcome the limited volume rates of standard OCT and two-photon angiography, we used a custom-built Bessel beam two photon microscope to monitor a 713×713×120 μm volume at 0.57 Hz for instances of stalled capillary flow. This was more than 4 times faster than OCT angiography of a similar volume. The increased speed led to a bottleneck in analysis, which was previously done fully manually. We semi-automatically extracted intensity along the centerline of all capillaries of interest and used it to flag time points of interest in all capillaries. The high false positive rate meant that our correlation metric was not sufficiently robust to directly detect stalls. But it was enough to reduce the time burden of analysis by about half (from 12 hours to ~6 hours), while identifying the vast majority of stalling events. The latter is especially important as the results of the fully manual analysis indicated that observers missed as many as 2/3 of all the stalling events. This is understandable given the complexity and size of even a single dataset, and given that it is easier to confirm that the automatically identified stalling events are truly a stalling event as opposed to the fully manual approach which requires inspection of all capillaries at all time points.

While we were able to greatly reduce the analysis burden while improving accuracy, stalling analysis was still time consuming. Deep learning-based tools have shown promise in distinguishing these events from normal flowing capillaries^26^, which could further reduce or eliminate the need for human observers. Due to the increased rate of stall detection with our approach compared with traditional two photon microscopy, the relatively small number of datasets used in our studies still contain processed results from over 6,000 capillaries that were monitored for 10 minutes each. Therefore, a sufficiently large training set of stalled and flowing capillaries can be created with fewer datasets compared to previous acquisition methods^26^. Our analysis would also likely benefit from improved and automated extraction of capillary traces from the Bessel beam data^26–28^.

We also leveraged the ability of Bessel beam to monitor pial artery dynamics simultaneously with capillary stalling (Figure 5a). As has been shown by others, vessel diameter correlates highly with vessel intensity^22^(Figure 5b), as a larger volume of intravascular dye is excited when the vessel diameter is increased. This provides an easy to calculate metric for vessel diameter, which we used to observe drops in stalling point prevalence around large dilation events (Figure 5c), as stalls were presumably cleared due to the transient increase in flow and perfusion pressure. This is consistent with recent work showing that environmental enrichment reduced the number of capillaries plugged by injected microspheres^29^. But here we show that there is on average fewer stalled capillaries around larger arterial dilation events *in vivo*, demonstrating the potential for dynamic reductions in stalling from spontaneous flow changes, not just under direct functional activation^13^. This effect was shown with large and isolated dilation events, but more sophisticated analysis can be done to better understand the relationship between stalling and neurovascular coupling, or even stalling and resting state oscillations.

To demonstrate the ability of our approach to detect changes in stall characteristics, we compared the statistics between young and old mice. In the older group, two mice underwent a photothrombosis model of stroke and were reimaged 1 week post-stroke. At baseline, the older mice had more than twice the number of capillaries that stalled at any given time (point prevalence) and in total (incidence) but had a larger share of stalling events that were shorter in duration (Figure 4f) resulting in a lower cumulative stall duration. This is further confirmed when removing shorter stalling events from analysis, resulting in a stall incidence similar to the younger group. The stall incidence of longer stalling events was consistent with that reported with OCT^13^, whose slower frame rate would not be sensitive to shorter events. Stroke resulted in a significant rise in stalling point prevalence and cumulative stall duration. This indicates that there is an increase in the duration of stalling events, even in the contralesional hemisphere. When recomputing the stall statistics using only longer events, an increase in incidence is also observed (Supplemental Figure 1). Circulation of neutrophils can still be elevated at this 1 week stage in recovery^30^, which could drive increases in stalling across the cerebral vascular network, including in the contra-lesional hemisphere. However, a larger scale study would be needed to confirm these phenomena. Future work can also address the source of longer vs shorter duration stalls using fluorescent labeling methods. For example, neutrophils adhering to the vessel wall could be the main driver of longer stalls. This would be consistent with previous work, where administering Anti-Ly6G to interfere with the adhesion of neutrophils reduced stalling^14,18^.The question of whether neutrophils primarily drive longer stalls could be addressed with multi-color imaging using Rhodamine 6G as a second fluorophore ^12,14^ (Supplemental Movie 2).

The ability to detect shorter stalls also raises the question of the shortest stall duration that should be considered during analysis. Previous work was only sensitive to longer stalls due to the nature of the measurement approaches and showed that the reduction in long stalls was critical and increased total cerebral blood flow well in agreement with a computational model^14^. In particular, it was found that a decrease in stalls from 1.8% to 0.72% resulted in a 13% increase in total cerebral blood flow^14^ Further, modeling of capillary transit times has shown that overall flow heterogeneity in the capillary network reduced oxygen delivery efficiency, even for the same total cerebral blood flow^19,31^, and we expect that increased stalling also leads to increased intracapillary flow heterogeneity. There is also predicted upstream and downstream effects of flow stoppages such that even brief stalls could theoretically increase spatial flow heterogeneity^32^, meaning flow heterogeneity is created across the local network. The increase in short stalls we found in our aged mice would be consistent with the increased spatial capillary speed heterogeneity found in older mice, where pockets of hypoxic tissue were also found more frequently^33^. On the other hand, while brief stalls are likely driving increases in flow heterogeneity and resultant pockets of hypoxic tissue, the diffusion of oxygen from the capillary tissue acts as a low pass filter of capillary oxygen dynamics^34^, and smooths out brief drops in oxygen, such that brief stalls would not likely induce critical levels of hypoxia in the adjacent tissue.

Since the imaging volume rate is equivalent to the imaging frame rate with Bessel beam two photon microscopy, it offers the potential for imaging capillaries at very high volume rates^22^. The frame rate used in our study was relatively slow compared to what is achievable with even standard linear galvanometers, but was used to allow for large field of view mages with a high signal to noise ratio. Further work needs to be done to understand what duration of stall is required to significantly affect oxygen delivery and to determine the optimal temporal resolution to capture their dynamics. Regardless, future investigations into capillary stalling will greatly benefit from high speed, high throughput measurements of capillary stalling statistics. Bessel beam two photon microscopy offers an ideal platform with an improved volume rate over conventional approaches and multicolor imaging of fluorescently labeled cells and plasma that can easily monitor hundreds of capillaries to study these events.

## Methods

### Animal Preparation

All procedures were reviewed and approved by the Boston University Institutional Care and Use Committee (IACUC). Animals were implanted with a custom titanium headpost and either a full crystal or two half crystal skulls (LabMaker), as described previously^35,36^. Animals were allowed to recover from surgery for three weeks before undergoing acclimation and imaging. During acclimation to head fixation, animals were head fixed for increasingly longer durations, and given sweetened condensed milk as a treat. Animals were monitored for signs of distress and removed early as necessary to avoid excessive stress. Acclimation sessions started at 15 minutes and continued daily until animals were comfortable with head fixation for 90 minutes.

### Two photon Microscopy

All imaging was performed under a home-built Bessel beam two photon microscope with a 100 μm axial PSF. An axicon based Bessel module was used to generate an annular illumination focused at the back pupil plane of the objective (Olympus 25x, 1.0 NA). Microscope control and data acquisition was done using Scanimage ^37^(Vidrio Technologies). Fluorescence was excited at 920 nm with a tunable Ti:Sapphire laser (Insight X3, Spectra Physics) with an electro-optic modulator (Conoptics 305-105-20) for fast power control. The excitation and emission paths were split by a primary dichroic (Semrock DI03-R785) and fluorescence was collected and detected by two cooled PMTs (Hamamatsu H74422-40 and −50). The fluorescence was split by a secondary dichroic (Semrock FF555-DI03) and passed through either a red (Semrock FF01-607/70) or green (Semrock FF01-502/60) before being detected by the PMTs.

### In vivo Imaging

Animals were briefly anesthetized and retro-orbitally injected with 30-50 μL of 150 kDa FITC-Dextran or Texas Red-Dextran (5% in PBS), and then head-fixed in a custom rig. The rig was then mounted on a motorized stage setup under the microscope. The custom rig used a goniometer stage for control of tip and tilt control on the headbar and craniotomy. The stage was adjusted such that the desired region of interest was perpendicular to the imaging plane of the microscope. Survey raster scans were used to check field flatness and match region of interests across sessions before acquisition of full times series. 512×512 pixel images of a 713×713×120 μm volume of view was scanned at 0.57 Hz for approximately 10 minutes (350 total images) and saved for offline processing. During the imaging sessions, the mouse was checked every ~30 minutes for signs of distress and given a treat of sweetened condense milk like in training sessions.

### Data Processing

Time series data was saved as tif series directly from ScanImage and used for analysis. For manual processing, data was loaded into a custom MATLAB GUI similar to that used for OCT angiogram stall analysis ^13,18^. Using a zoom feature, a sub-ROI was focused on for analysis. The series was then inspected frame by frame looking for instances of stalling. Stalls were classified as frames where all red blood cell (RBC) shadows were stationary between frames. Any movement in the RBCs or change in the number of visible RBCs was classified as flowing (non-stalled). Analysis was completed for each sub-ROI until all capillaries were observed. Once all stalled capillaries had been found, each stall was then analyzed frame by frame and all frames where the capillary stalled were manually annotated. The manually annotation as well as segmented capillary and centerline (see below) were then saved for group analysis. Finally, the total number of capillaries in the ROI was counted and recorded. Only capillaries with clear RBC contrast were counted.

For correlation-based detection, capillary centerlines were semi-automatically extracted by first marking all capillaries of interest with a single click on each capillary. Capillaries were selected based on having sufficient contrast to see RBC shadows along the length of the vessel. To extract centerlines, images first underwent vessel enhancement to increase the contrast of the thinner capillaries using the Frangi vesselness filter function fibermetric in MATLAB. Images were then thresholded for a black and white image of vessels versus background and then a skeleton was extracted using the bwskel function. The skeleton was converted to a graph following the approach used by Xiang et al.^23^ The nearest edge segment to each manual click was then used at the capillary centerline. The intensity along the centerline was then extracted frame by frame and then used to generate the length vs time (LT) images similar to a kymograph. To reduce the effects of motion on the LT images, an image was cropped around each segment and then locally registered. This transform was then applied to the centerline to keep it centered on the vessel even during and after motion.

### Vessel Diameter

To estimate vessel diameter, 20 lines spaced 2 μm apart were drawn along the width of the vessel. The resulting intensity profiles were then averaged and normalized. The diameter was estimated to be the full width at half max (FWHM) of the resulting profile. Fluorescent intensity was measured using a small rectangular ROI drawn inside the same vessel, and mean intensity was used to calculate ΔF/F.

Large dilation events were identified manually from 9 of the 10 ROIs from the validation dataset after extraction of vessel dynamics. One ROI was excluded because it lacked any significant vasodilation events. To prevent the potential confound of successive dilation events, we only analyzed isolated dilation events. Events where another large vasodilation occurred within 10 frames (17.5 seconds) were excluded.

### Validation Experiment

Validation was performed on 10 datasets that had already underwent fully manual analysis. Capillaries were selected as described above and LT plots generated. Capillary segments of 5 pixels or less were excluded from analysis. For each time point (row) in the image, the intensity was correlated with the intensity of the next frame (row), resulting in the frame-to-frame vessel intensity correlation over time. This correlation trace was thresholded, and all suprathreshold time points were flagged as stalls. A new observer (different from the observer who conducted fully manual analysis) then inspected every flagged result and marked it as a true or false positive. The new observer also inspected stalls marked during fully manual analysis, including those not flagged by the correlation metric. The analysis was restricted to only flagged capillaries and time points, to minimize the time burden of analysis.

### Photothrombotic Stroke

Stroke was induced following a vessel-targeted photothrombotic model^24^ to provide a clinically relevant model. The stroke was performed under a custom built intrinsic optical signal imaging (IOSI) and laser speckle contrast imaging (LSCI) system that allowed for continuous flow monitoring during occlusion.

The location of the stroke was determined using spatial frequency domain imaging (SFDI)^25^ taken on Day 2 and on Day 7 in a short session prior to Bessel imaging. ROIs at baseline were selected based on the region the target vessel fed and were predicted to be close to the stroke boundary. One ROI was also taken in the contralateral hemisphere to serve as a control.

## Supplemental Figures

**Supplemental Figure 1.**
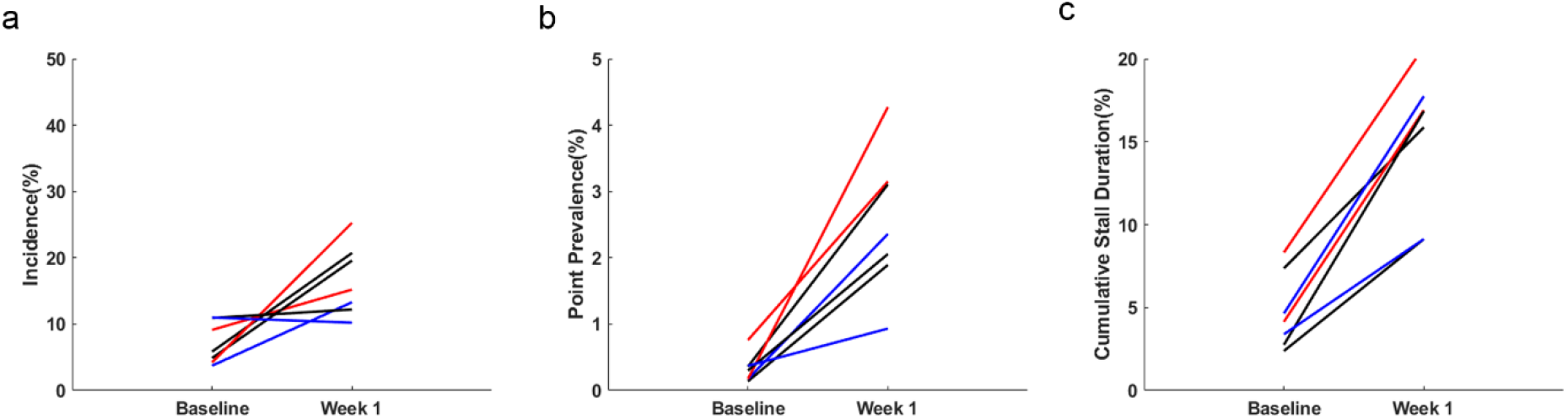
Stall statistics at baseline and one week post stroke calculated using only longer duration stalling events. Colors indicate region relative to the stroke (red indicates stroke core, black peri-infarct, and blue contralesional hemisphere) (a) Stall incidence (b) Stall point prevalence (c) Cumulative stall duration

## Notes

### Competing Interest Statement

The authors have declared no competing interest.

